# Spatial and Temporal Consistency of Brain Networks for different Multi-Echo fMRI Combination Methods

**DOI:** 10.1101/2021.08.18.456877

**Authors:** J. Pilmeyer, G. Hadjigeorgiou, R. Lamerichs, M. Breeuwer, A.P. Aldenkamp, S. Zinger

**Affiliations:** Department of Electrical Engineering, Eindhoven University of Technology, Eindhoven, the Netherlands; Department of Biomedical Engineering, Eindhoven University of Technology, Eindhoven, the Netherlands; Department of Research and Development, Epilepsy Centre Kempenhaeghe, Heeze, the Netherlands; Department of Neurology and Clinical Neurophysiology, Maastricht University Medical Center, Maastricht, the Netherlands; Philips Research, Eindhoven, the Netherlands; Philips Healthcare, Best, the Netherlands

## Abstract

The application of multi-echo functional magnetic resonance imaging (fMRI) studies has considerably increased in the last decade due to its superior BOLD sensitivity compared to single-echo fMRI. Various methods have been developed that combine the fMRI time-series derived at different echo times to improve the data quality. Here we evaluated three multi-echo combination schemes, i.e. ‘optimal combination’ (T_2_*-weighted), temporal Signal-to-Noise Ratio (tSNR) weighted, and temporal Contrast-to-Noise Ratio (tCNR) weighted combination. For the first time, the effect of these multi-echo combinations on functional resting-state networks was assessed in the temporal and spatial domain, and compared to networks derived from the second echo (35 ms) functional images. Sixteen healthy volunteers were scanned during a 5 minutes resting-state fMRI session. After obtaining the networks, several temporal and spatial metrics were calculated for their time-series and spatial maps. Our results showed that, compared to the second echo network time-series, the Pearson correlation and root mean square error were the most consistent for the optimal combination time-series and the least with those derived from tSNR-weighted combination. The frequency analysis further suggested that the time-series from the tSNR-weighted combination method reduced hardware- and physiological-related artifacts as reflected by the reduced power for the associated frequencies in almost all networks. Moreover, the spatial stability and extent of the networks significantly increased after multi-echo combination, primarily for the optimal combination, followed by the tSNR-weighted combination. The performance of the tCNR-weighted combination lacked robustness and instead varied remarkedly between resting-state networks in both the temporal and spatial domain. The results highlight the benefits of multi-echo sequences on resting-state networks as well as the importance of adjusting the choice of multi-echo combination method to the research question and domain of interest.

## Introduction

Over the last decades, functional magnetic resonance imaging (fMRI) has provided numerous novel insights into brain activation patterns of healthy individuals and patients with neurologic disorders. The application of fMRI has proven to be useful in research domains such as classification of healthy versus diseased individuals [1], the prediction of optimal treatment [2, 3] or identification of biologically-based subtypes of neurologic disorders [4] and also in clinical domains such as the planning of brain tumor surgery and localization of epileptogenic lesions [5].

In addition to the measurement of brain activity when the subject is performing a task or is presented with stimuli (task-based), one can study the activity of the brain when the subject is ‘at rest’, i.e. lies still in the MRI scanner (resting-state fMRI). Independent component analysis (ICA) is a widely applied method for the analysis of such resting-state fMRI data. By assuming that the data consist of linear mixtures of unknown independent variables and by maximizing their non-Gaussianity, ICA facilitates the identification of these statistically independent components. These components can be thought of as distinct functional brain networks, so called resting-state networks (RSNs). RSNs have been widely studied in the healthy population and neurologic disorders. For example, the default mode network (DMN), which has been found to be active during self-reflection and inactive during attention-demanding tasks [6], can be robustly detected during resting-state fMRI and hyperactivation of this network has been linked with major depressive disorder. Other consistently identified RSNs are the executive control network (ECN), the salience network (SN) and the visual network (VN).

In standard (i.e. single-echo) fMRI, a single slice is acquired after each radiofrequency excitation pulse at a specific echo time (TE). Despite its non-invasiveness and high spatial resolution, single-echo fMRI is prone to different noise sources that can be of physiological, motion, thermal or hardware-related origin [7]. To increase the sensitivity of the acquired blood oxygenation level-dependent (BOLD) contrast in fMRI and to prevent signal dropout, an MRI technique called multi-echo (ME-)fMRI has been developed [8]. In ME-fMRI, slices are acquired at different TEs following each RF-pulse. This allows for the estimation of *T_2_\**, a time constant which is inversely related to the decay of the transverse magnetization and reflects the tissue properties and magnetic field inhomogeneities, for each voxel and time point [9]. The acquisition of signals of voxels with different *T_2_\** decay rates results in images with different signal intensities and tissue contrasts. The signal acquired at each *TE* can be estimated using a mono-exponential fit:

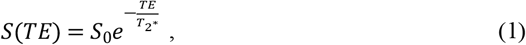

where *S(TE)* is the signal acquired at echo time *TE* and *S_0_* is the signal acquired at *t* = 0. From this equation it can be seen that signals acquired at smaller *TE*s have higher signal intensity than longer *TE*s. However, the contrast between gray and white matter and cerebral spinal fluid has been shown to be higher at longer *TE*s [9]. Thus, by combining the images that are acquired at different *TE*s, additional physiological information can be obtained, thereby increasing the sensitivity of the BOLD contrast. Several of such multi-echo combination methods have been developed. Examples are the temporal Signal-to-Noise Ratio (tSNR) combination [10] and the T_2_*-weighted combination (also called ‘optimally combined’, OC) [8], the latter takes into account the variation of T_2_* – known to be dependent on the brain location and tissue type [11] – over the brain.

There is a scarcity of studies in which the effect of the different ME combination methods on fMRI data is investigated. Previous research compared the BOLD sensitivity of multi-echo fMRI data that were either combined by a simple echo summation scheme, a temporal Contrast-to-Noise Ratio (tCNR or alternatively called ‘PAID’ method) weighted combination or the previously described T_2_*-weighted combination to data acquired by single-echo fMRI. They found significant sensitivity increases of more than 7% and 11% for the T_2_*-weighted and tCNR-weighted combination method, respectively. They demonstrated even larger increases in regions with a very short or long T_2_* that are more susceptible to signal dropout in single-echo fMRI [12]. Another recent study showed that the T_2_*-weighted and the tCNR-weighted combination methods increased the tSNR of fMRI time-series by more than 30% over the whole brain compared to single-echo. These combinations also increased the temporal contrast-to-noise ratio of fMRI time-series in emotional and finger tapping tasks [13]. Controversially, results from a dual-echo study indicated that there were no significant sensitivity advantages on a group-level by using tSNR-weighted or tCNR-weighted multi-echo combination over simple multi-echo averaging [10].

The current paper presents the first research of the evaluation of the effect of different echo combinations on resting-state networks. In our approach, three different echo combination methods will be applied to multi-echo resting-state fMRI data: tSNR, tCNR and OC echo combination. The temporal and spatial consistency between resting-state networks resulting from the different echo combinations will be compared between each other and between the second echo (SE) reference. Based on previous work we expect large temporal and spatial RSN differences between the multi-echo combination methods and the second echo reference. Additionally, we hypothesize that the type of echo combination has different effects on either the temporal or spatial properties of the RSNs.

## Materials and Methods

### Subjects

Sixteen healthy subjects participated in this research and all of them gave informed consent. None of the subjects had a medical history of neurologic or psychiatric disorders and they were between 20-65 years old (mean age = 43.4 years old, 11 females and 5 males). The study was approved by the Academic Center for Epilepsy Kempenhaeghe (Heeze, the Netherlands) based on METC approval N16.098.

The following sections describe the main steps from the acquisition of the MRI data up until the evaluation of the consistency metrics between different echo combination methods. Fig 1 shows a schematic of these analysis steps required to obtain the final results.

**Fig 1.**
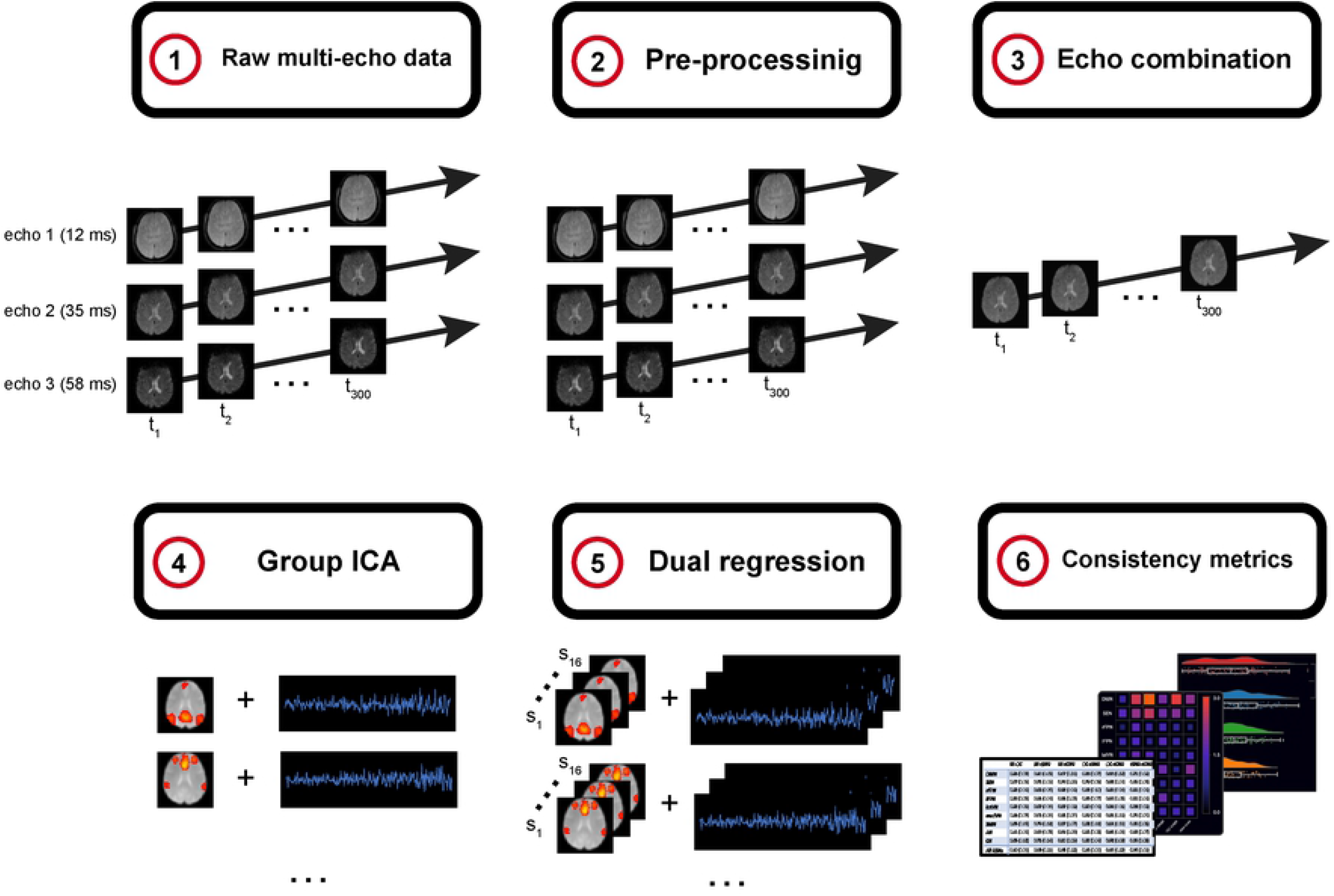
Schematic of the required analysis steps to obtain the consistency metrics between resting-state networks. 1: The raw data consists of multi-echo data, i.e. 3 time-series acquired at different echo times. 2: Pre-processing steps are applied. 3: The three time-series are combined using the different combination methods, resulting in a single combined time-series. For the single-echo reference, the second echo time-series is taken. 4: Group-ICA results in groupwise spatial maps and time-series. 5: Individual spatial maps and time-series are obtained from dual regression. 6: The RSNs of all subjects and combination methods are evaluated for temporal and spatial (in)consistencies.

### Data acquisition

Scanning was performed on a Philips Achieva MRI scanner (3 Tesla). At first, a T1-weighted anatomical scan was recorded using a 3D spoiled gradient-echo sequence (repetition time (TR) = 8.3 ms, TE = 3.5 ms) resulting in a matrix size of 240 × 240 × 180 with isotropic voxels of 1 mm^3^. Multi-echo (3 echoes) images (300 volumes per echo) were acquired using a gradient-echo EPI sequence (TR = 2000 ms, TE = 12 ms, echo spacing = 23 ms). In total, 26 slices with a slice thickness of 4.5 mm were obtained with an in-plane resolution of 3.5 × 3.5 mm and a final in-plane resolution of 3.5 × 3.5 mm. A SENSE acceleration factor of 2.7 was applied in the read-out direction.

### Image pre-processing

Data were pre-processed with fMRIPrep [14]. Each T1-weighted image was corrected for intensity non-uniformity and was skull-stripped, both implemented in ANTs [15, 16]. For the functional images, a reference volume and its skull-stripped version were generated using a custom methodology of fMRIPrep. The fMRI images were then slice-time corrected using AFNI [17] and corrected for head-motion by registration to the reference image. Furthermore, the fMRI time-series were high pass filtered (>0.01 Hz), and voxel-wise variance normalized whereas each volume was spatially smoothed with a σ = 7 mm Gaussian kernel, all carried out in FSL [18].

### Multi-echo combination methods

Following the pre-processing of each separate echo, the echoes were combined according to different echo combination schemes. Three following multi-echo combination algorithms were selected for comparing resting-state network consistency.

#### Optimally combined (T_2_*-weighted)

The T_2_*-weighted multi-echo combination method, usually referred to as “optimally combined” (OC), was developed by Posse et al. [8] and is the combination scheme implemented in the multi-echo denoising method ME-ICA by Kundu et al. [19]. The OC algorithm optimizes contrast by estimation of *T_2_\** for each voxel [8], which can lead to reduction of susceptibility artifacts and thermal noise [20]. The weights for echo combination are calculated as follows:

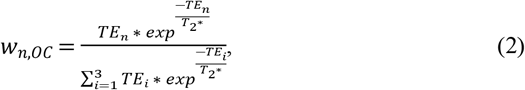

where *TE* is the echo time and *T_2_\** is the estimated T_2_* from each voxel. The indices *n* and *i* reflect the echo number with the difference that *i* is used to sum over the three different echoes for normalization of the weights. After calculating the weights for all three echo images, the combined fMRI time-series of each voxel can be calculated by taking the weighted average using the optimally combined weights.

#### tSNR-weighted

In the tSNR-weighted echo combination method, first the *tSNR* of every voxel’s time-series of each echo image is calculated. The *tSNR* is defined as the mean a time-series divided by its standard deviation. Subsequently, the following equation can be used to calculate the weight, *w_n,tSNR_*, of the image with echo *n* [10]:

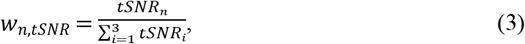

where *tSNR_n_* is the temporal signal-to-noise ratio of the image with echo *n*, and *i* the echo index used to sum over all three echo images. After calculating the weights for all three echo images, the combined fMRI time-series of each voxel can be calculated by taking the weighted average using the tSNR-based weights.

#### tCNR-weighted (tSNR- and TE-weighted)

The tCNR-weighted echo combination approach, introduced by Poser et al. [12], combines the *TE* and *tSNR* values for each echo image reflecting the temporal contrast-to-noise ratio of the images. The advantage of this method is that it does not make any assumptions about the signal and noise because it is measured from the data while it simultaneously incorporates the echo time simultaneously. The *tCNR* weights are calculated as follows:

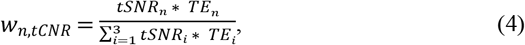

Again, the weighted average is calculated to retrieve the final tCNR-weighted combined fMRI image.

#### Second echo reference

The resting-state networks that are extracted from the multi-echo combined fMRI images will be compared to those derived from the second echo fMRI images. The corresponding echo time of the second echo is equal to 35 ms.

### Network extraction and selection

Each functional image in native space was coregistered to its corresponding T1w-image using a 6 degrees-of-freedom (rotation and translation) boundary-based registration, implemented in FSL’s FLIRT. Then, functional images in T1w-space were spatially normalized to MNI-space by a linear 12 degrees-of-freedom (rotation, translation, scaling and skewing) registration resulting in an isotropic resampling resolution of 3.5 mm. Resting-state networks (RSNs) were extracted by group-level independent component analysis (ICA) using FSL’s MELODIC [18]. The number of spatially independent components was set to 30 in order to obtain isolated RSNs with minimal contamination from other networks or subdivision into smaller components [21]. All of the subjects’ fMRI time-series were temporally concatenated before identification of independent components. After estimation of the group-average RSNs, a two-step dual regression procedure was performed in FSL [22] to obtain individual RSN time-series and spatial maps. The first step resulted in individual time courses by regression of the group-average spatial maps into each subject-specific 4D dataset. The individual spatial maps were obtained after the second step during which regression of the time courses into the 4D dataset of the first step was applied.

Networks were identified using a goodness-of-fit approach with the RSN atlas from Smith et al. as reference [23]. From this RSN atlas, a mask image of each resting-state network was used and compared to each of the 30 independent components obtained from ICA. The goodness-of-fit approach was applied to each comparison and was defined as the average z-score of the IC voxels that fall within the mask image subtracted from the average z-score of the IC voxels that lie outside of the mask image. The network that scored the highest on this goodness-of-fit score was selected for each IC. Subsequently, a visual inspection was performed to check the quality of the match. One of the ICs did not match any of the IC templates but partly overlapped with the executive network. In addition to the executive network, the involvement of the salience network (including the anterior cingulate cortex and anterior insula) was apparent in that IC. Therefore, this network will be referred to as the salience-executive network (SEN) [24].

In the end, the following nine resting-state networks were identified: the default mode network (DMN), salience-executive network, right and left frontal parietal network (rFPN/lFPN), lateral and medial visual network (latVN/medVN), somatosensory network (SMN), auditory network (AN) and cerebellar network (CN). Fig 2 shows the general identified networks [23, 24] whereas S1 Fig presents the networks derived after each multi-echo combination method and the second echo reference.

**Fig 2.**
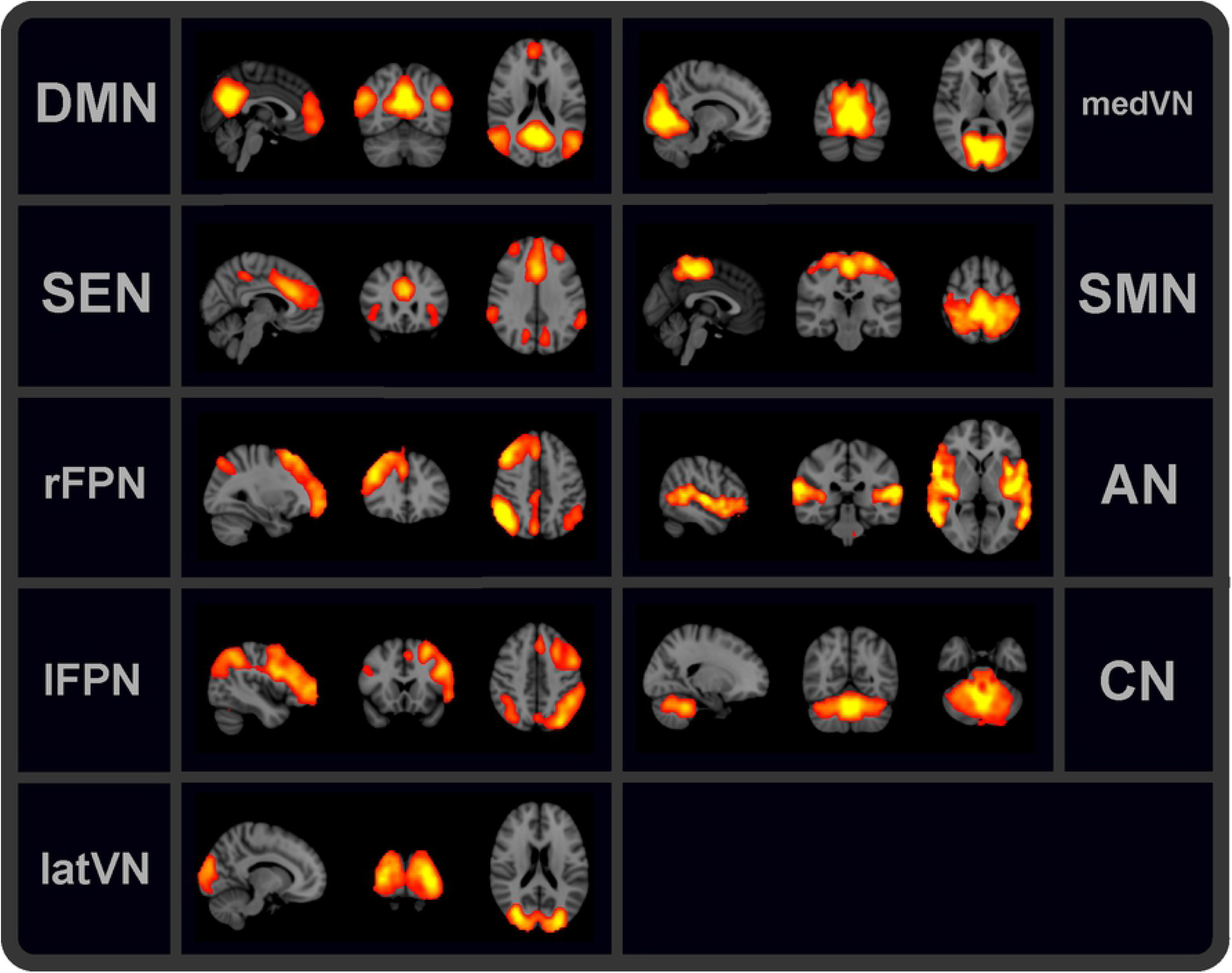
Identified resting-state networks. Nine common resting-state networks were identified from the fMRI data following independent component analysis.

### Consistency metrics

Multiple metrics were calculated to compare the properties of the RSNs derived from the combination methods described above. These metrics provide insights into the spatial, i.e. related to the RSN maps, and temporal, i.e. related to the RSN time-series, domain.

#### Temporal consistency metrics

The first temporal consistency metric was the *Pearson correlation*, which was implemented to evaluate the synchrony between two time-series. The second metric that was applied is the *root mean square error* (RMSE) which penalizes the differences in values of two time-series and is therefore dependent on the absolute values of both two time-series. This is opposed to the Pearson correlation, which is based on differences of relative values. Finally, in order to get insights of the spectral properties of the fMRI time-series, Welch’s *power spectrum density* (PSD) was calculated, as implemented in the Python-based package Nitime [25]. The power spectrum reflects the contribution of each frequency to the signal and could therefore give an indication of differences in BOLD fluctuations or noise components between the echo combination methods.

#### Spatial consistency metrics

The first spatial consistency metric was the total *number of ‘active’ voxels*. The number of ‘active’ voxels is defined as the amount of voxels that exceed a z-score of 3 [26]. The total count of the number of active voxels was used to assess the spatial extent of the RSNs [26]. In addition, the *maximum z-score* for each network was obtained. The maximum z-score of each network gives an indication of the stability of each RSN [26]. The last spatial consistency metric that was used for evaluation is the *Dice coefficient* [27]. The Dice coefficient estimates the amount of overlap between the spatial maps of the RSNs.

## Results

### Temporal consistency

The overall Pearson correlation was relatively high, with no correlations below 0.66 (Fig 3A or S1 Table). Compared to the second echo reference, the OC time-series were surprisingly similar with a correlation of 0.9 ± 0.08 over all RSNs whereas the tCNR-weighted time-series were the least correlated with a Pearson correlation of 0.83 ± 0.11. Focusing on individual networks, the tSNR-weighted and tCNR-weighted combination methods showed lower correlations (within a range of 0.71 – 0.80) for the SEN and CN time-series compared to the SE time-series. Interestingly, the DMN time-series was the least correlated of all networks for the three tCNR-weighted comparisons (within a range of 0.66 – 0.71). This is possibly due to the inclusion of more anterior voxels for the tCNR-combined RSN (S1 Fig) leading to less correlated time-series compared to the other echo combination methods.

**Fig 3.**
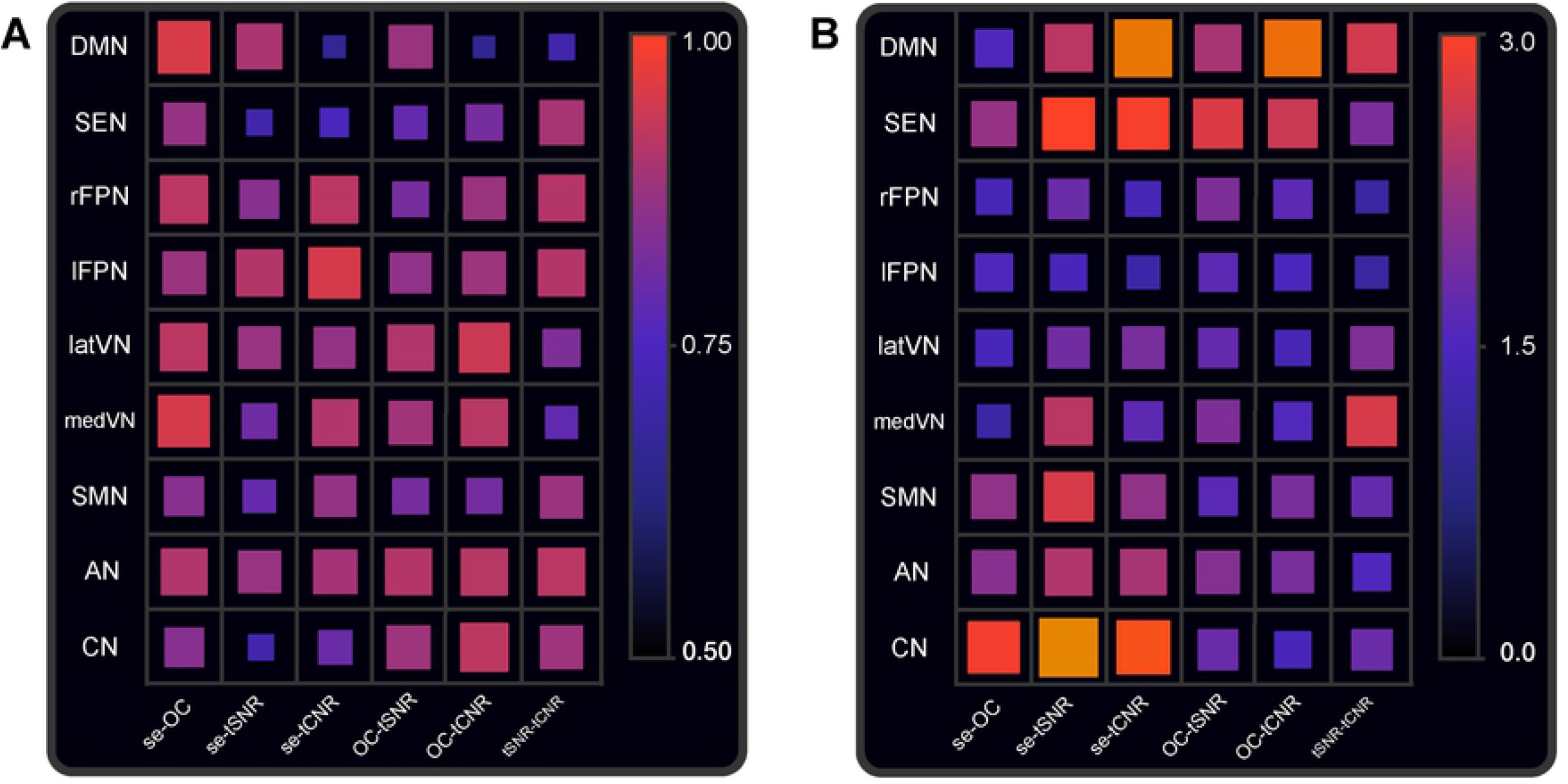
Average Pearson correlation and root mean square error between time-series for each RSN and multi-echo combination method and second-echo reference. (A) Average Pearson correlation between time-series of RSNs (y-axis) for different multi-echo combination methods and the second echo reference (x-axis). The elements on the x-axis represent each possible comparison between the multi-echo combination methods and the second echo reference. The average over subjects was taken. The size of the squares as well as the colour-coding reflect the value of the correlation. (B) The root mean square error is shown with the same axes as Fig 2A The size of the squares as well as the colour-coding reflect the value of the RMSE.

From Fig 3B and S2 Table it can be seen that the RMSE results were mostly in agreement with the results from the Pearson correlation, showing low RMSE values for networks with high correlations. Similar to the Pearson correlation, OC differed the least from the SE reference, followed by tCNR and tSNR (RMSE of 1.84 ± 1.00, 2.29 ± 1.09 and 2.46 ± 1.26 over all RSNs for the three SE-comparisons, respectively). On the contrary, the RMSE plots revealed that the time-series of the DMN, SMN and AN for tSNR and tCNR also deviate considerably from the second echo in terms of absolute values (RMSE range from 2.19 – 3.00).

In the frequency range between 0 and 0.05 Hz the SE time-series showed the highest power for almost all RSNs, followed by OC (Fig 4). Exceptions on this observation were the SEN and SMN for which the power of the tCNR series dominated. Noteworthy, the power of the tSNR was consistently the lowest for this frequency range among RSNs. After 0.05 Hz, for all combination methods and RSNs, the power considerably decreased up until 0.2 Hz. From 0.2 Hz and higher, the power increased again but the trend was less unambiguous than in the frequency range between 0 and 0.05 Hz. For the DMN, SEN and the rFPN, the power of the OC time-series dominated, peaking around 0.23 Hz. The tCNR had the highest power for both visual networks whereas the SE had the highest power contribution for the other RSNs in this frequency range.

**Fig 4.**
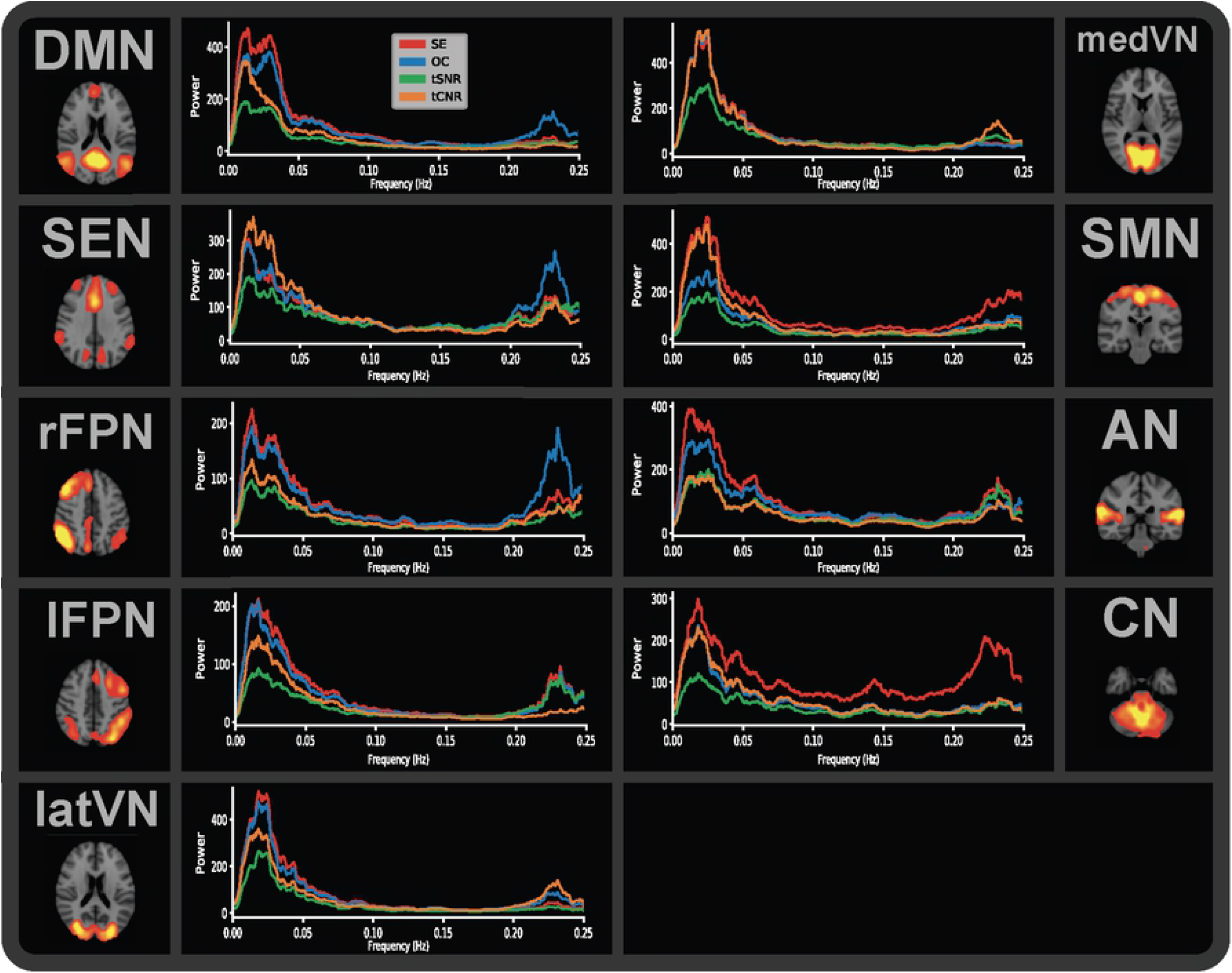
Frequency plots of the time-series for each RSN and multi-echo combination method and second echo reference. Welch’s power spectrum density of the time-series of each RSN and for each multi-echo combination method as well as the second echo reference.

### Spatial consistency

Distribution plots of the maximum z-score for all RSNs and subjects indicated an improvement of network stability for OC or tSNR-weighted multi-echo combination as the maximum z-score for these combination methods is significantly higher than the second echo z-scores (p = 2.9 * 10^−3^ and 4.6*10^−2^, respectively), see Fig 5A. Moreover, the peak z-score for the OC and tSNR combination method was significantly higher than the tCNR method (p = 3.5*10^−6^ and p = 1.6^−4^, respectively). The amount of ‘active’ voxels, i.e. with a z-score larger than 3, significantly increased when using ME over SE for all three combination methods (p = 1.3 * 10^−6^, p = 3.1 * 10^−6^ and p = 4.7 * 10^−5^ for OC, tSNR and tCNR, respectively), see Fig 5B.

**Fig 5.**
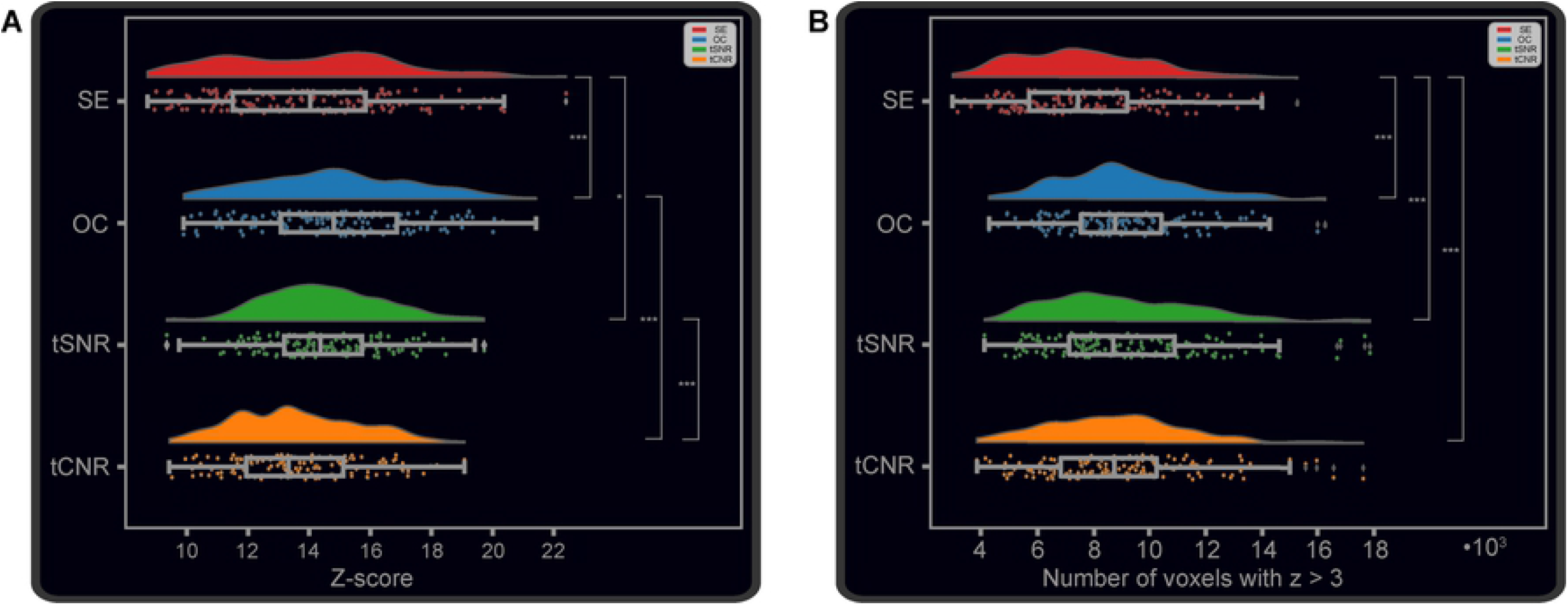
Distribution plots of the maximum z-score and total count of voxels with a z-score > 3 per multi-echo combination method. (A) Distribution plots of the maximum z-score per multi-echo combination method, reflecting spatial stability. Each data point represents one of the nine resting-state networks of a single subject. (B) Distribution plots of the total count of voxels with a z-score > 3 per multi-echo combination method, reflecting spatial extent. * = p-value < 0.05, ** = p-value < 0.01, *** = p-value < 0.001.

Finally, the Dice coefficient values were mostly in the range between 0.25 – 0.75. From Fig 6 and S3 Table it can be seen that the RSNs from OC were again the most similar in comparison to the RSNs from the SE data. This was in agreement with the Pearson correlation and RMSE results. Overall, the networks that showed the least overlap with the SE RSNs were the SEN, medVN and the CN. In addition, the DMN from the tCNR-weighted combination showed to have very little overlap with the RSNs from the other methods, which is probably due to the more prominent anterior part of the DMN. The most consistent networks in terms of overlap can be found when comparing OC with tSNR-weighted combination for which all Dice coefficient values are higher than 0.50.

**Fig 6.**
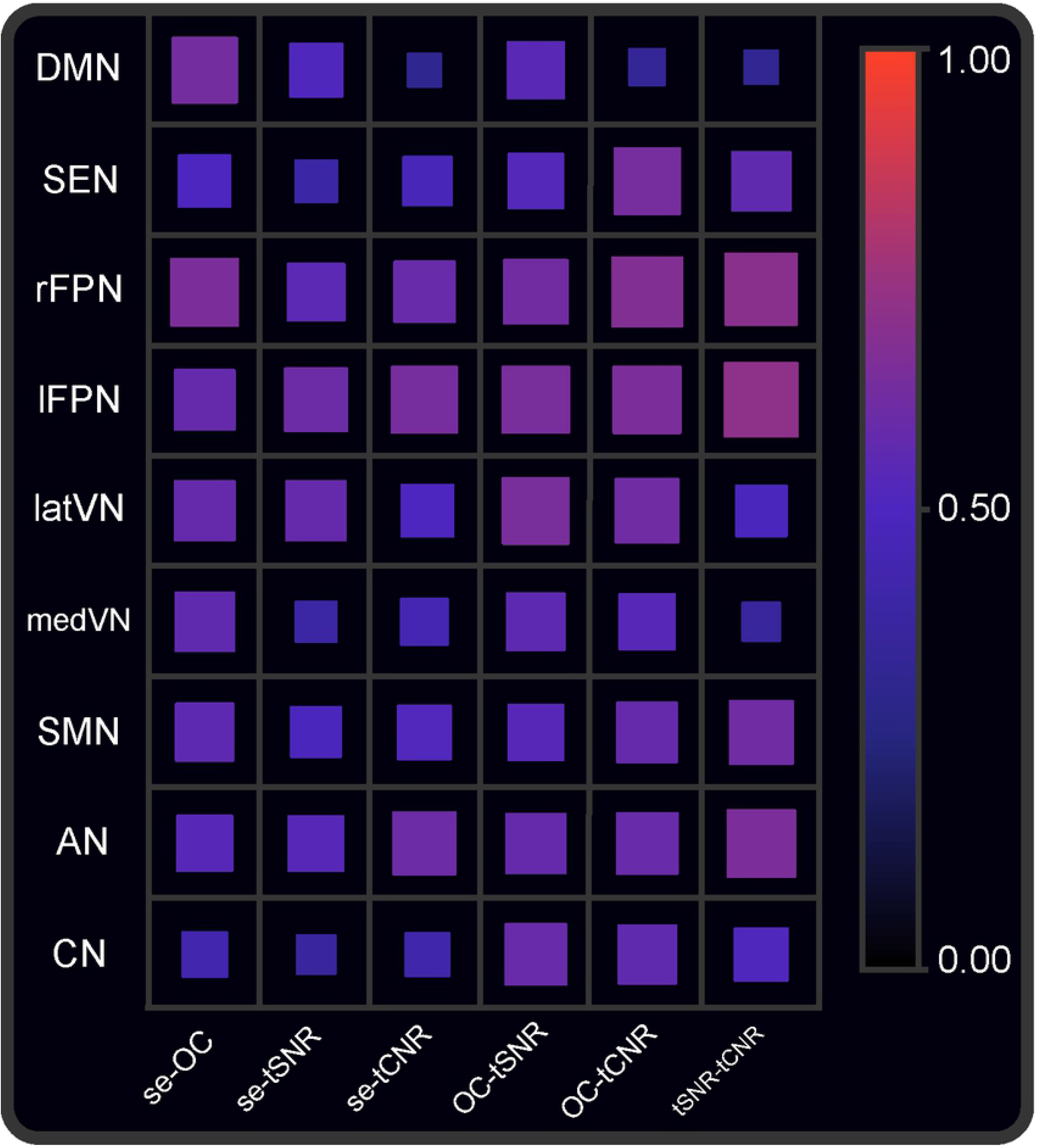
Dice coefficient for each RSN and multi-echo combination method and second echo reference. Average Dice coefficient values for each RSN between spatial maps of different multi-echo combination methods as well as the second echo reference. The average over subjects was calculated.

## Discussion

In this paper we evaluated the consistency of the spatial and temporal properties of RSNs between different multi-echo combination methods. It was shown that the choice of ME combination method results in substantially varying RSNs in terms of time-series correlation, RMSE and frequency spectra, as well as spatial stability, extent and overlap.

### Second echo

The most remarkable finding of network comparison with the second echo reference was the high temporal inconsistency with the tSNR-weighted combination and higher spatial stability and extent for OC.

The tSNR-weighted combined RSN time-series deviated the most from the SE time-series compared to the other multi-echo combinations. The lower power of the time-series from the tCNR and especially the tSNR in the frequency ranges between 0 and 0.05 Hz and 0.2 and 0.25 Hz likely caused the overall reduction of correlation and increase in RMSE. Interestingly, these frequency ranges are often associated with noise. The 0.2 – 0.25 Hz range is linked with contamination of respiratory signals while the 0 – 0.05 Hz is frequently obscured by scanner drift (0 – 0.01 Hz) and respiration-induced CO_2_ fluctuations (0 – 0.05 Hz) [7]. Assuming most of the power in these ranges comprises noise sources, ME combination reduces its amount with tSNR being the most effective, followed by tCNR and OC. Yet, from these results it is uncertain whether true BOLD signals in the 0 – 0.05 Hz range are also affected. More structural removal of these artifacts, e.g. by RETROICOR [28] or ME-ICA [19, 20], or simulations could aid in the process of identifying the origin of this power reduction.

The OC spatial maps appeared to benefit the most from multi-echo combination with regard to the network stability and extent. One possible explanation is that the BOLD contrast, as observed in the spatial maps, is closer to optimal for OC due to its voxel-specific T_2_*. Previous research has already shown that T_2_* is dependent on the location in the brain [11]. Moreover, dropout of signals in tissue with T_2_* values that are more deviating from the TE, can be more prevented in OC combination compared to tSNR- or tCNR-weighted combination [20]. The increase in BOLD contrast and minimization of signal dropout could have led to the increase in the maximum z-value and number of ‘active’ voxels. The temporal metrics and Dice coefficient of the OC networks were highly consistent with the OC method, suggesting that the OC combination method exclusively improves the spatial network properties.

The effect of multi-echo combination in the temporal domain was mostly evident in the SEN and CN and in lesser proportions the DMN, SMN and AN as all ME combination methods show relatively low correlation and high RMSE values in these networks. Possibly, the coverage of more voxels in distinctive brain areas as a result of ME combination could have produced these temporal differences. For example, the spatial maps of the SEN (S1 Fig) contained a more prominent cluster of voxels in the supramarginal gyri (the posterior lateral clusters) and the posterior cingulate cortex (the two medial, most posterior clusters) while the CN voxels covered a larger area of the cerebellum. This observation was also reflected in the lower Dice coefficients for these networks. Different neuronal physiological properties of those areas could have been the underlying reason for the differences in the concerned RSN time-series and corresponding frequency spectra between SE and ME.

### Optimal combination

As discussed before, the most noticeable finding for the OC method was its higher spatial network stability and extent compared to SE. On the other hand, the OC RSNs showed similar temporal metrics and Dice coefficient values as SE. More specifically, the spatial maps of most RSNs for OC are highly overlapping with an exception of the SEN, CN and to a lesser extent the AN compared to the second echo reference. The power spectrum for most of the RSNs for OC was slightly lower in the 0 – 0.05 Hz and 0.2 – 0.25 Hz ranges, of which the latter is associated with respiratory artifacts. Increases of BOLD sensitivity using the OC method have been observed before [12, 13], suggesting that the power decrease could be related to the reduction of artifacts or increases of the contribution of the neuronal signal. Nonetheless, this does not explain the increases of power in the 0.2 – 0.25 Hz domain for the DMN, SEN, rFPN and latVN. Possibly, the OC is more robust at reducing low frequency noise components in time-series of RSNs and less so at removing frequencies above 0.2 Hz. Additional removal of artifactual IC components from the time-series, e.g. by detection of and regressing out the noise-related components as implemented by ME-ICA [19, 20], could help to determine whether these higher frequency components > 0.2 Hz in the OC RSN time-series are indeed of non-neural origin.

From the results, it can be concluded that the OC method might be the preferred choice in research related to the identification of functional maps rather than evaluating the network temporal aspects. It could be questioned whether the increased peak z-scores and number of active voxels are not originating from noise sources. If this were the case, however, the power spectrum in the noise frequency ranges would presumably be considerably higher for OC compared to SE as noise sources often obscure the true neuronal-related activity [29]. Likewise, the power in the frequency range 0.05 – 0.15 Hz, which is usually associated with true neuronal BOLD signals due to the physiological nature of the BOLD signal and the hemodynamic response [29], is mostly equal for the OC time-series in comparison with SE.

### Temporal Signal-to-Noise Ratio combination

The RSN time-series derived after tSNR-weighted combination showed the least consistency with the SE reference. Overall, the power spectrum of the outer frequency ranges was the lowest of all combination methods in almost all RSNs while the middle frequency ranges around 0.05 - 0.2 Hz were still preserved. Spatially, the tSNR-weighted combined maps approach the OC maps, as indicated by the high Dice coefficients. Despite their compactness, the distribution plots and p-values of the network stability and extent show slightly lower, yet insignificant, values compared to the OC method. Nevertheless, the tSNR-weighted combination method showed significantly increased peak z-scores compared to SE and tCNR-weighted combination. Combined, tSNR-weighted combination had the highest impact on the temporal and spatial features of the RSNs. The spectral analysis hints at tSNR-weighted combination being the most RSN improving ME combination method temporally. This can be explained by the fact that tSNR is a temporal-based metric that attributes the heaviest weights to echo images with the highest signal-to-noise ratio. This could have led to the observed noise reduction. Research related to the investigation of the temporal effects of RSNs might opt to combine multi-echo fMRI data using the tSNR-based combination. An example of such a study could be the evaluation of RSNs using the wavelet coherence analysis in which phase shifts are a key factor [30]. Artifacts, such as the respiratory artifacts, that are left in the RSN time-series could bias these analyses significantly as these are often emerging as repeating temporal patterns. Studies investigating the temporal aspects of RSNs might benefit from the tSNR-based ME combination due the power reduction in the frequency ranges of these physiological confounders.

### Temporal Contrast-to-Noise Ratio combination

Overall, the tCNR-weighted combination showed inconsistent results between networks. In terms of temporal consistency to the SE reference, the tCNR-weighted combination showed up in between the OC and tSNR-weighted combination methods. The correlation and RMSE metrics showed to have a high network-dependency as some networks were highly consistent with SE while others were relatively inconsistent. This trend was also noticeable from the power spectrum in which some networks, such as the rFPN and AN, had low power while others such as the SEN and medVN showed the highest power of all ME combination methods and even the SE reference. Spatially, the RSNs were largely overlapping with the OC RSNs but lacked network stability and extent, making the potential multi-echo benefit to be neglectable for the tCNR-weighted combination.

### Alternatives

One of the most popular multi-echo fMRI denoising methods is ‘ME-ICA’ by Kundu et al. [19, 20]. This technique first combines the echoes by a weighted average using the OC weights, after which ICA is applied. Based on the IC’s TE-dependence, ICs are either classified as BOLD or non-BOLD components. Subsequently, the time-series from the non-BOLD ICs are regressed out from the fMRI time-series to clean up the data. The ME-ICA algorithm has shown to be robust, minimizing motion and other physiological artifacts as well as preserving the DMN’s functional connectivity [19, 31]. Moreover, it was found that ME-ICA reduced the power in the higher frequency domains of ten common RSNs from Smith et al. [23] while preserving the frequencies of spontaneous BOLD fluctuations [31]. Despite its promise on improving the spatial and temporal properties of the RSNs, the ME-ICA denoising method was out of the scope of this study as the goal was to evaluate the inconsistencies between combination methods without additional denoising.

A novel method called T_2_*FIT-weighted combination was recently introduced by Heunis et al. [13] in which the identical weights as the OC method are used but with the difference that T_2_* maps are fitted per time point instead of per time-series. This resulted in a significant gain in tSNR of ~37% over the whole brain compared to single-echo fMRI, as well as larger effect sizes and functional contrasts. To put things in perspective, the tSNR of the time-series from OC and tCNR combination increased by ~31% and ~32%. Future studies could investigate whether these results from the T_2_*FIT-weighted combination translate into more robust RSNs temporally and spatially.

## Conclusions

The application of multi-echo fMRI in future studies is warranted thanks to its significant increase in BOLD sensitivity when compared to conventional single-echo fMRI. A limited amount of studies has compared the effect of different echo combination strategies on BOLD sensitivity and signal-to-noise ratio. Here we evaluated, for the first time, the effect of different echo combination methods on resting-state networks. To achieve this, we analyzed the temporal and spatial properties of common resting-state networks derived from ICA and also compared them to networks derived from the second echo. We found that the time-series from the tSNR-weighted combination were the least consistent compared to the second echo networks. This was accompanied with decreased power for almost all RSNs in the frequency ranges that are associated with hardware- and physiological-related artifacts. The time-series from the optimal combination were consistent with those derived from the second echo reference but the spatial maps achieved the highest scores on spatial stability and extent. The performance of the tCNR-weighted combination lacked robustness and instead varied remarkedly between RSNs. The results highlight the benefits of multi-echo sequences on resting-state networks as well as the importance of adjusting the choice of multi-echo combination method to the research question and domain of interest. Limitations of the study include the relatively high repetition time and low sample size. Future studies could investigate the effect of the promising T2*FIT-weighted combination and optimal combination with additional denoising (ME-ICA) on resting-state networks.

## Acknowledgments

We would like to express our gratitude to Dr. Ir. Bernas (Donders Institute for Brain, Cognition and Behavior) for our fruitful discussions and for the insights provided by his PhD thesis. It helped us to formulate the research questions, investigated in this paper.

## Supporting information captions

**S1 Fig. Identified resting-state networks for each multi-echo combination method and second echo reference.** Nine common RSNs were identified from the fMRI data for each multi-echo combination and the single-echo method. The maps were thresholded at a z-score > 3. Abbreviations: SE = single echo, OC = optimally combined, tSNR = temporal signal-to-noise ratio, tCNR = temporal contrast-to-noise ratio, DMN = default mode network, SEN = salience-executive network, rFPN/lFPN = right/left frontoparietal network, lat/mVN = lateral/medial visual network, SMN = sensorimotor network, AN = auditory network and CN = cerebellar network.

**S1 Table. Pearson correlation of fMRI time-series between different multi-echo combination methods and the single-echo reference per RSN.**

The mean and standard deviation (in between brackets) is shown. Abbreviations: SE = second echo, OC = optimally combined, tSNR = temporal signal-to-noise ratio, tCNR = temporal contrast-to-noise ratio, DMN = default mode network, SEN = salience-executive network, rFPN/lFPN = right/left frontoparietal network, lat/medVN = lateral/medial visual network, SMN = sensorimotor network, AN = auditory network and CN = cerebellar network.

**S2 Table. Root mean square error of fMRI time-series between different multi-echo combination methods and the single-echo reference per RSN.**

The mean and standard deviation (in between brackets) is shown. Abbreviations: SE = second echo, OC = optimally combined, tSNR = temporal signal-to-noise ratio, tCNR = temporal contrast-to-noise ratio, DMN = default mode network, SEN = salience-executive network, rFPN/lFPN = right/left frontoparietal network, lat/medVN = lateral/medial visual network, SMN = sensorimotor network, AN = auditory network and CN = cerebellar network.

**S3 Table S3. Dice coefficient of spatial maps between different multi-echo combination methods and the single-echo reference per RSN.**

The mean and standard deviation (in between brackets) is shown. Abbreviations: SE = second echo, OC = optimally combined, tSNR = temporal signal-to-noise ratio, tCNR = temporal contrast-to-noise ratio, DMN = default mode network, SEN = salience-executive network, rFPN/lFPN = right/left frontoparietal network, lat/medVN = lateral/medial visual network, SMN = sensorimotor network, AN = auditory network and CN = cerebellar network.

